# Modelling Competition for Nutrients between Microbial Populations Growing on a Solid Agar Surface

**DOI:** 10.1101/086835

**Authors:** Boocock Daniel, Conor Lawless

## Abstract

**Motivation:** Growth rate is a major component of the evolutionary fitness of microbial organisms and an excellent surrogate for cell health. In the high-throughput procedures QFA and SGA, microbial cultures are inoculated in an array on solid agar, and growth is measured to obtain quantitative fitness estimates. Neighbouring cultures, which are often different genetic strains, consume nutrients at different rates, creating gradients in nutrient density. We believe that diffusion of nutrients between cultures is affecting growth in these experiments. Current analysis does not account for this; instead, it is assumed that cultures grow independently. I use a network model of nutrient diffusion and nutrient-dependent growth, to correct for competition, to try to improve the accuracy and precision of fitness estimates. I test the model against QFA data from studies on telomere function in *Saccharomyces cerevisiae*. Ultimately, the model might be used to improve the reliability of screens for genetic interaction and drug sensitivity.

**Results:** I fit the competition model to a QFA plate from Addinall et al. (2011). Using far fewer parameters (387 vs 1152), the new model fits timecourses with similar closeness to the previous model. Fitness estimates are less precise for the fastest growing strains, but more precise for the majority of strains (36 out of 50). Fitness rankings agree in the positions of the fastest and slowest growing strains, but disagree in the middle positions. In a cross-plate validation experiment, the competition model overestimates timecourses to a similar degree that the previous model underestimates. A different method of fitting is required to find globally optimal solutions which might improve reliability.

**Availability and Implementation:** CANS, a Python package developed for the analysis in this paper, is freely available at https://github.com/lwlss/CANS.

## 1 INTRODUCTION

The bacterium *Eschericia coli* and yeast *Saccharomyces cerevisiae* are unicellular organisms studied as a model prokaryote and eukaryote respectively. Much of their genomes have been conserved in other species over billions of years of evolution (O’Brien *et al*., 2005). *S. cerevisiae* was the first eukaryote to have its entire genome sequenced (Goffeau *et al*., 1996), and is commonly used as a basis for the study of other eukaryotes including human cells.

Bacteria and yeasts grow in colonies. In favourable conditions, growth is exponential, and growth rate is a major component of fitness; faster growing strains quickly come to dominate populations. At a certain point, growth becomes limited and a stationary phase is reached, so pressure also exists to use resources efficiently. In short, fitness is governed by competing pressures on growth rate and yield (Dethlefsen and Schmidt, 2007). It is possible to observe growth to estimate fitness (Baryshnikova *et al*., 2010a; Addinall *et al*., 2011). Cell cultures are grown on either the surface of a nutrient rich solid agar, or suspended in a liquid mixture containing nutrients, and cell density is measured over time. Fitness is defined by measures such as growth rate. Identical strains might grow differently between mediums causing fitness estimates to differ (Baryshnikova *et al*., 2010a). Here, I focus on estimating the fitness of microbes grown on a solid agar surface.

Fitness estimates can be used to infer genetic interaction or drug response. Using high-throughput methods, this can be conducted on a genome-wide scale (Costanzo *et al*., 2010; Andrew *et al*., 2013). In a typical genetic interaction screen, a strain is created with a mutation in a query gene (background). Second deletions are introduced to create different double mutants. Genetic interactions are inferred by comparing the growth of a double mutant to the growth of the two corresponding single mutants. If the double mutant is fitter than predicted given the observations of single mutant fitness, then the deletion is said to suppress the defect of the query mutation. If the double mutant is less fit than predicted given the observations of single mutant fitness, then the deletion is said to enhance the defect of the query mutation. Either scenario suggests that the two genes interact. Due to redundancy, single deletions are often non–lethal. This allowed Costanzo *et al*. (2010) to explore genetic interactions for ~75% of the *S. cerevisiae* genome.

Synthetic Genetic Array (SGA) and Quantitative Fitness Analysis (QFA) are high-throughput fitness-screening methods, which use solid agar, and make quantitative estimates of microbial fitness (Baryshnikova *et al*., 2010b; Banks *et al*., 2012). In SGA, cultures are pinned onto plates. Typically, in QFA, dilute liquid cultures are inoculated onto plates. In both procedures, cultures are arranged in rectangular arrays. Each culture is a single strain and each plate my may contain 100s or 1000s of different strains. Whole genomes can be explored using different combinations of query genes and deletions. I study QFA, in which a model is fit to growth curves, and fitness estimates are defined in terms of model parameters. Typical QFA procedures use 16×24 arrays (384 spots). In contrast, SGA procedures use a single endpoint assay of culture area to quantify growth, and, typically, use larger 32x48 arrays (1536 dots). In QFA, inoculum density may be varied to capture more or less of the growth curve; the most dilute cultures are inoculated with ~100 starting cells (Addinall *et al*., 2011). Plates are incubated at a controlled temperature and periodically removed for whole plate photographs to be taken. Incubation lasts several days, to capture both the exponential and stationary growth phases. The software Colonyzer (Lawless *et al*., 2010) converts optical density measurements in photographs to cell density timecourses for each culture. Past analysis fit the logistic growth model (1). The differential form and solution of the logistic model (Verhulst, 1845) are given in (1), where *C* is cell density, *r* is a growth constant, and *K* is a carrying capacity.

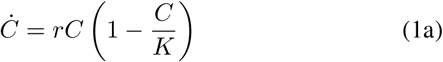

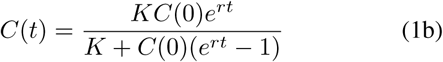

The logistic model is a simple mechanistic model describing self-limiting growth and has a sigmoidal solution. Growth begins exponentially, with rate *rC*, and curtails as the population size increases and cells begin to compete for some limited resource. Cell density reaches a final carrying capacity, *K*, at the stationary phase. In QFA, nutrients must diffuse through agar to reach cells growing on the surface. *K* might be reached when nutrients run out, or growth becomes diffusion limited and is approximately stationary. Fitting the logistic model to a 384 culture plate, requires culture level parameters for *r* and *K*, and either plate or culture level parameters for *C*(0), making a total of 769 or 1152 parameters.

The growth constant *r* could be used as a fitness measure. Addinall *et al*. (2011), however, define a more complicated fitness measure as the product of Maximum Doubling Rate (MDR) and Maximum Doubling Potential (MDP), which they calculate from logistic model parameters. MDR is the doubling rate at the beginning of the exponential growth phase. MDP is the average number of cell divisions between inoculation and stationary phase.

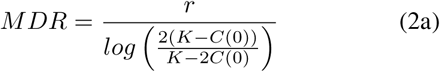

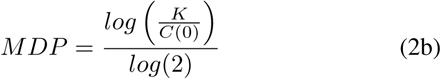

To improve the quality of fits, QFA now uses the generalised logistic model which requires an extra shape parameter for each culture (Banks *et al*., 2012). Standard and generalised logistic model r are not equivalent, so comparison relies on MDR and MDP as fitness measures. Either model can be fit using the QFA R package (Lawless *et al*., 2016).

Since QFA aims to differentiate strains by fitness, many different strains are grown on one plate, and fast and slow growing strains often grow side-by-side. This is exemplified in Figure 1, which shows a section of a QFA plate from Addinall *et al*. (2011). Cultures were inoculated with approximately equal cell density, but grew to different sizes, at different rates. In timecourses from this plate, all cultures appear to reach the stationary phase at a similar timepoint, despite starting with the same amount of nutrients and growing at different rates. This suggests a global growth-limiting effect, which might be caused by an interaction between cultures. I test whether the interaction is a competition effect, caused by diffusion of nutrients along gradients formed between fast and slow growing neighbours. Competition has implications for growth estimates; growth will appear faster or slower for each neighbour, than it would if cultures grew independently. The experiment in Figure 2 provides further evidence of an interaction between neighbours. Identical strains were repeated in the same positions on two plates, but on one plate every second column was left empty. Cultures in Figure 2a, with fewer neighbours, grew larger than the same cultures in Figure 2b.

**Figure 1:**
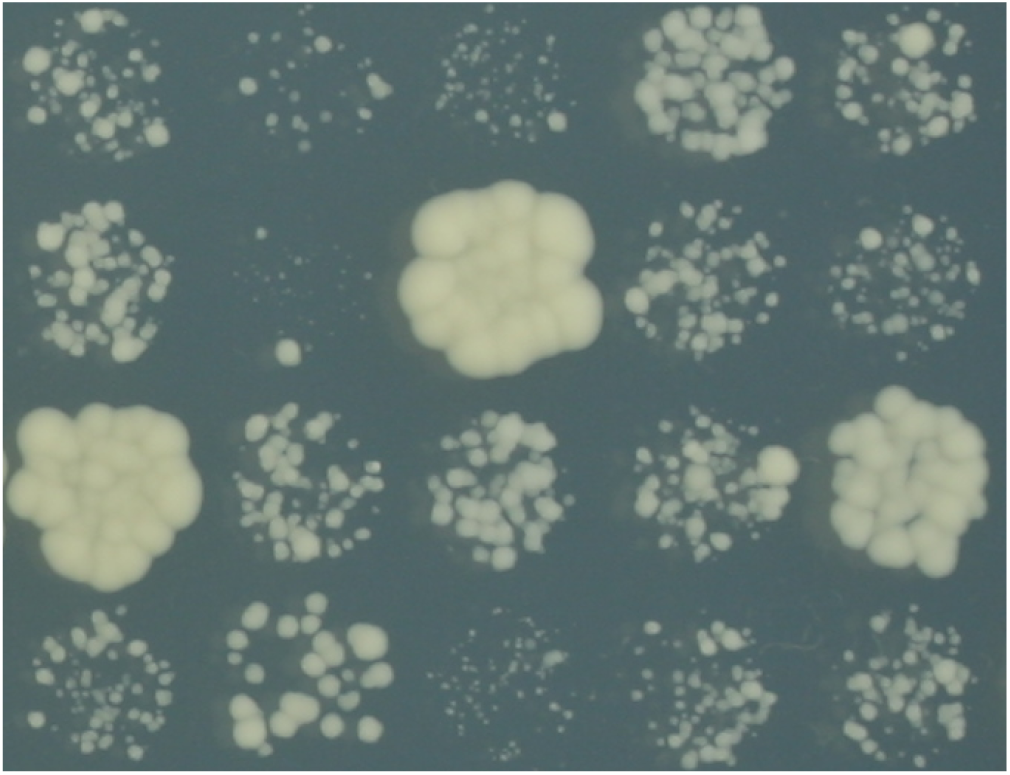
4×5 section of a QFA plate (P15). Cropped from a 16×24 format solid agar plate inoculated with dilute S. cerevisiae cultures. Image captured after ~2.5 days incubation at 27° C.

**Figure 2:**
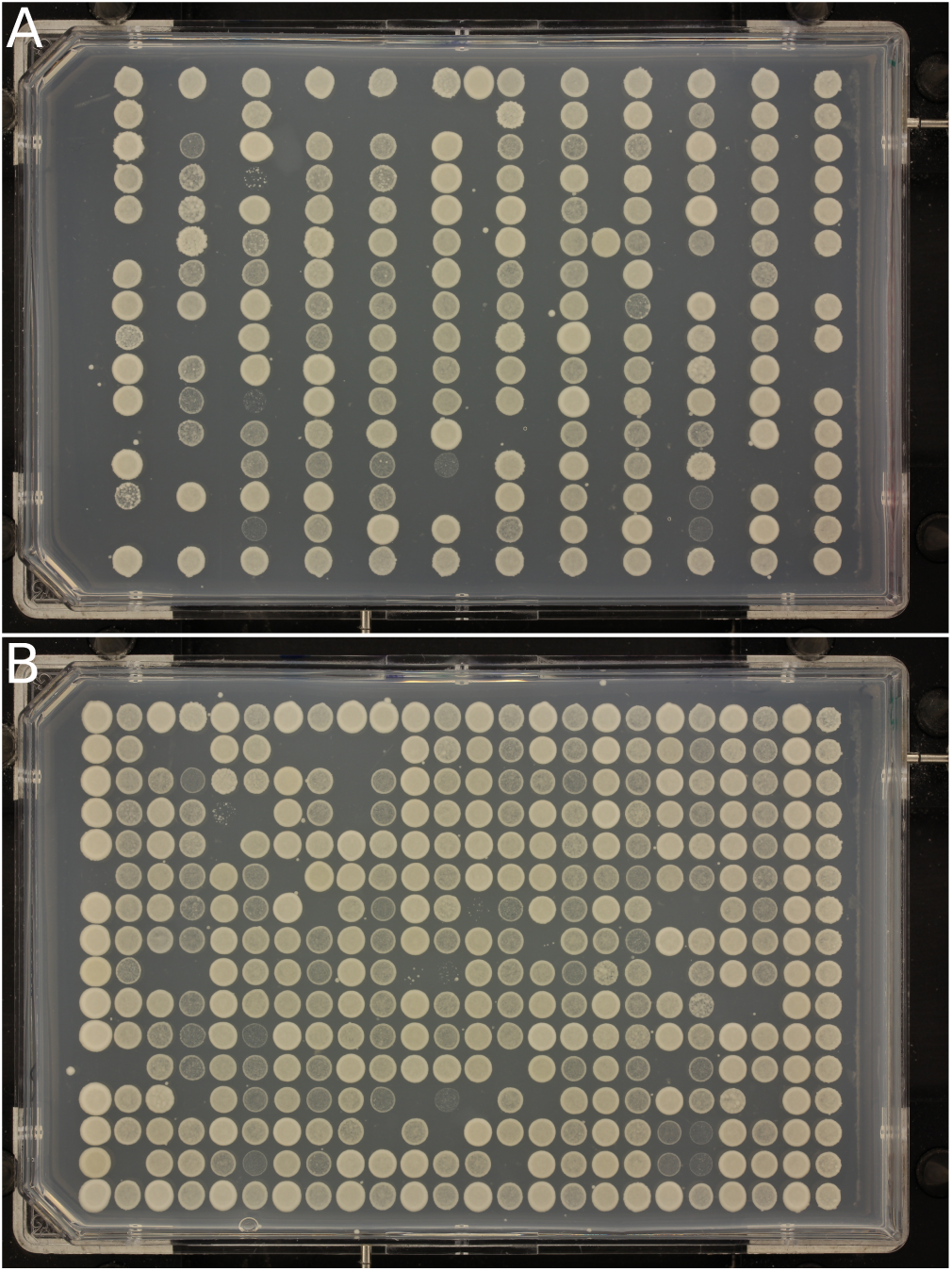
Plates from a QFA experiment designed to examine competition. **A)** “Stripes”; **B)** “Filled”. Identical strains are repeated in the same positions on each plate, except in every second column in A, where locations are left empty. The extra columns in B are a transposition of the columns to the left, but with a different query mutation. The leftmost column in B contains repeats of a control with a neutral deletion. Cultures were inoculated with a higher cell density than in Figure 1.

Current QFA analysis, using the logistic model, assumes that cultures grow independently. A sigmoidal curve fits QFA data poorly in many cases. I aim to correct for competition using a network model of nutrient diffusion and nutrient-dependent growth, to improve the fit of individual growth curves, and increase the accuracy and precision of fitness estimates.

There have been attempts to reduce and correct for competition effects experimentally and statistically. This does not require explicit knowledge or modelling of the source of interaction. For example, QFA data for edge cultures is usually discarded because of noise, but Addinall *et al*. (2011) grow repeats of a neutral deletion in edge locations, rather than leave them empty, because of concerns about competition. Plates could also be repeated with randomised culture positions. However, this is unlikely to remove all bias; for instance, the fastest growing strains would always have slower growing neighbours. In an SGA study, Baryshnikova *et al*. (2010a) use various statistical techniques to improve the correlation between repeats of the same strain in different positions. I expect that an explicit model of competition, fit to whole growth curves, will provide a better correction and improve fitness estimates, without requiring extra repeats. Furthermore, a model might identify and explain the source of competition. Simulations of a successful model could be used, not only to analyse existing data, but to compare experimental designs, and predict ways to reduce competition experimentally. Poisoning of cultures by a signal molecule, such as ethanol, which *S. cerevisiae* produces in the metabolism of sugars by fermentation, is another possible source of interaction. QFA does not measure nutrients or signal, so, if more than one type of interaction is significant, it will be very difficult to fit a model. In such circumstances, experimental and statistical methods might be the best approach.

Reo and Korolev (2014) use a 2D diffusion equation model to simulate nutrient dependent growth of a single bacterial culture on a pertri dish. They create a sink for nutrients from culture growth, and equate the flux of nutrients through culture area with the rate of increase in culture size. They keep culture density constant and allow culture area to vary.

This model could be adapted for QFA, if instead, culture area is approximated as constant and culture density is allowed to vary. It would, however, be too computationally intensive to fit such a model to a full QFA plate in three-dimensions, especially if the model is to be used to analyse many plates from high-throughput experiments. Therefore, a simpler model of nutrient diffusion is required.

We propose a network model of nutrient diffusion and nutrient-dependent growth (Figure 3), hereinafter the competition model, which uses reaction equations and a mass action kinetic approximation. Nutrient-dependent division of cells is represented by the reaction equation,

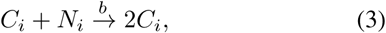

where *C* is a cell, *N* is the amount of some limiting nutrient required for one cell division, and *b* is a rate constant for the reaction. The identity of *N* is unknown, but possible candidates are sugar and nitrogen. Each culture, indexed i, has a separate reaction (3) with growth constant *b_i_*. Mass action kinetics gives the rate equation (4a) for the amount of cells, *C_i_*, in a culture, and the first term in the rate equation (4b) for the amount of nutrients, *N_i_*, associated with a culture.

**Figure 3:**
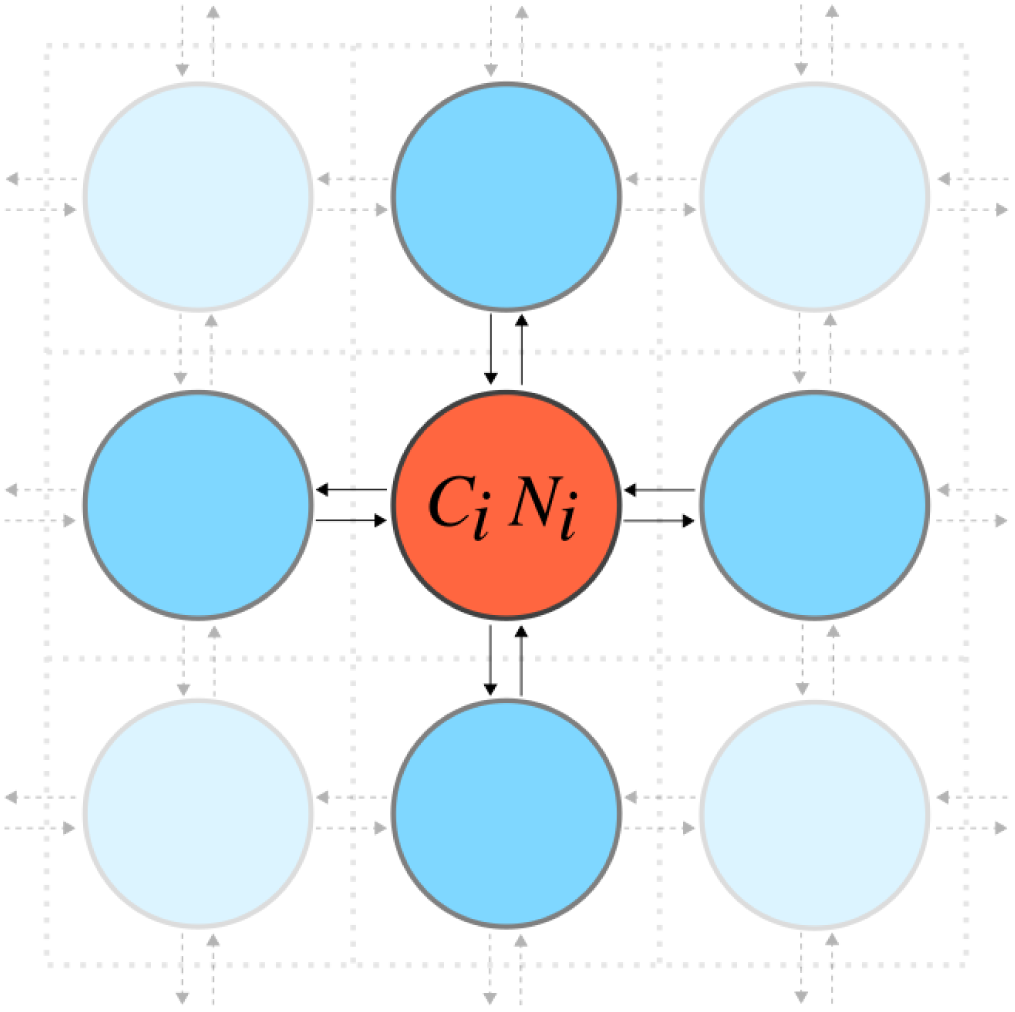
Schematic of the competition model. Cultures, represented by circles indexed i, grow in a rectangular array on the surface of a nutrient containing solid agar. The agar is divided into equal areas surrounding each culture. Each culture has an amount of cells, C_i_, and is associated with an amount of nutrients, N_i_. Arrows represent diffusion of nutrients between cultures. Darker blue circles, δ_i_, are the closest neighbours of the red culture, i.

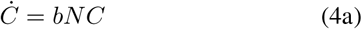

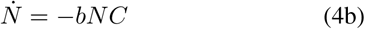

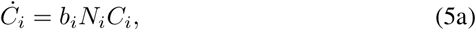

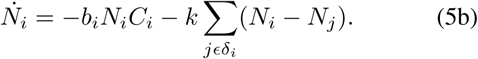

To arrive at the full competition model, the diffusion of nutrients along gradients between a culture and its closest neighbours, *δ_i_*, is modelled by reactions of the form,

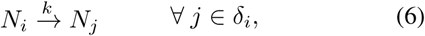

where *k* is a nutrient diffusion constant which is independent on culture location. Using mass action kinetics, the sum of diffusion reactions between a culture and its neighbours gives the second term in (4b).

Unlike the logistic model (1), the competition model has no analytical solution; instead, it must be solved numerically. When *k* = 0, the competition model is a special case of Monod kinetics (Monod, 1942) with an assumption of unsaturated nutrients and yield constant, the mass ratio of cells produced to nutrients consumed, equal to one. Considering conservation of mass, Kargi (2009) shows that in this case, the Monod model is equivalent to the logistic model and can be re-expressed as such, with substrate consumption implicit. We use the form with explicit substrate, hereinafter the mass-action logistic model, so that we can also model nutrient diffusion. Importantly, the equivalence of the competition and logistic models allows us to convert between parameters to compare fitness estimates (see Section 2.5).

In QFA, *C*_*i*_ are observed and *N*_*i*_ are hidden. Inoculum densities, *C*_*i*_(0), are often below detectable levels. When fitting the competition model, I assume that inoculum density is the same for all cultures, nutrients are evenly distributed throughout the agar at time zero, and k is constant across the plate. I can therefore infer *C*(0), *N*(0), and *k* at the plate level. Including growth constants, *b*_i_, for each of 384 cultures on a typical plate, makes 387 parameters in total. The competition model has roughly half as many parameters as the standard logistic model, and a third as many as the generalised logistic model.

The competition model makes many other approximations. It is a deterministic model which uses a continuum approximation for numbers of reactant species. In QFA, populations begin with ~100 cells, but quickly grow to reach thousands of cells, so this approximation appears valid. Mass action kinetics applies to reactions in a well stirred mixture, and is, perhaps, less valid for cultures growing on solid agar. A mass action approximation has, nevertheless, been successful in other situations where this assumption is questionable: in the Lotka-Volterra model of predator-prey dynamics (Berryman, 1992) and in signalling and reaction models inside cells (Aldridge *et al*., 2006; Chen *et al*., 2010). The order of a reaction also affects the rate equation, but the identity, and quantity of the nutrient molecule is unknown. Reactions also assume that all nutrients are converted to cells and include no model of metabolism. I justify the use of the competition model because in the independent limit it has the same solution as the logistic model, long used to model microbial growth. Furthermore, collectively fitting the competition model involves a large number of parameters and data points, and will require many simulations to be run. Computational feasibility necessitates the use of many approximations.

## 2 METHODS

### 2.1 CANS

I developed a Python package (CANS) for model composition, model simulation, parameter inference, and visualisation of results. I use CANS to fit the competition model to QFA data. CANS accepts cell density timecourses for locations in rectangular arrays of arbitrary size. SBML models are produced to document results of parameter inference, and for independent validation using other simulation tools. It is simple to create new models with reactions which are common to each location, including diffusion reactions between nearest neighbours. CANS is available at:https://github.com/lwlss/CANS.

### 2.2 The P15 dataset

I compared the logistic and competition model by fitting both to a single plate from a previous QFA study on *S. Cerevisiae* (Addinall *et al*., 2011). I call this plate P15. A 3x3 section is shown in Figure 1. P15 contains 384 cultures, all with background mutation *cdc 13-1*. *cdc13* is involved in telomere stability and *cdc13-1* is a temperature-dependent mutation. Each culture also has a deletion from a standard deletion library containing 49 different deletions thought to affect telomere function. There are 6 repeats of each of these deletions. There are also 14 repeats of a strain with a neutral deletion, *his*△. Edge cultures are inoculated with extra repeats of the neutral deletion, but results are discarded due to noise from reflections off plate walls. Inoculum density is ~100 cells, which is below the level of detection. P15 was incubated at 27° C, at which point *cdc13-1* starts to experience loss of function. Each culture has a cell density timecourse with 10 timepoints covering 4 days. Several features make P15 a good test case. Strains vary greatly in fitness, so competition should be present. Repeats of each strain increase statistical power. Furthermore, for several strains, independent spot tests have been published and may be used for validation (Maringele and Lydall, 2002; Zubko *et al*., 2004; Holstein et al., 2014; Foster *et al*., 2006).

### 2.3 The Stripes and Filled datasets

I used a two-plate QFA experiment (E. Holstein, personal communication, April 2016) to validate the competition model. Images of the plates, termed the “Stripes” and “Filled” plates, are shown in Figure 2. As for P15, the experiment uses strains of *S. Cerevisiae* with a background mutation, and a deletion in a gene relevant to telomere function. In contrast to P15, there are more deletions per plate and no repeats for most strains. Identical strains are repeated in the same positions on each plate, except in every second column in the Stripes plate, where locations are left empty. The common cultures have background mutation *cdc13-1*, as used in P15, and identical strains are inoculated from the same liquid culture to reduce genetic variation. The extra columns in the Filled plate are a transposition of the columns to the left, but with a different query mutation *rad75*Δ. The leftmost column in the Filled plate contains repeats of the neutral deletion, *his*Δ. Inoculum density is ~10,000 cells, which is much higher than that used in P15, and detectable at time zero. The plates were incubated together at 27° C, where the defect of *cdc13-1* is larger than that of *rad75*Δ. Each culture has a cell density timecourse with ~50 timepoints covering four days. This is about five times as many timepoints as collected for P15.

The different neighbour configurations cause identical strains to grow differently between the two plates. To validate the competition model, I calibrate parameters on the Filled plate and validate by simulating timecourses for the Stripes plate. This is possible because the Filled plate contains all of the strains present on the Stripes plate.

### 2.4 Solving and fitting

#### 2.4.1 Solving

CANS solves ODE models numerically using one of two packages: integrate from SciPy, and libRoadRunner. The method integrate.odeint solves ODEs written in Python, at user supplied timepoints. I used NumPy to vectorise competition model ODEs to optimise solving by this method. libRoadRunner is implemented in C++ and solves ODEs for models written in SBML. I access libRoadRunner through its Python API, and use the libSBML Python API to automatically generate SBML versions of the competition model for any size plate. For a 16×24 culture plate with cell density observations at 10 evenly spaced timepoints, libRoadRunner is approximately 10 times faster than SciPy’s integrate, solving in hundredths rather than tenths of a second. Unlike SciPys’s integrate.odeint, libRoadRunner solves only at evenly spaced timepoints. To fit P15, where each timecourse has 10 unevenly spaced timepoints, I simulated sequentially between adjacent timepoints. For the Stripes and Filled plates, which have ~50 unevenly spaced timepoints, I sampled 15 evenly spaced timepoints from a spline. I use SciPy’s interpolate.splrep to make 5th order B-splines with smoothing condition *s* = 1:0, and SciPy’s interpolate.splev to evaluate. The sampled timepoints range from time zero to the time of last observation. Solving then requires just one call to libRoadRunner’s RoadRunner.simulate. Using libRoadRunner on a modern CPU, fitting a 16×24 format plate with 10 unevenly spaced timepoints takes ~3 hours; fitting a full plate with 15 evenly spaced timepoints takes ~1 hour.

#### 2.4.2 Fitting the competition model

I processed raw QFA data using Colonyzer (Lawless *et al*., 2010;) to produce cell density timecourses. To these, I fit the competition model, using a gradient method, and normal model of measurement error, to make maximum likelihood estimates of parameters. The particular gradient method was a constrained minimisation: the L-BFGS-B algorithm from SciPy’s integrate package. I determined stopping criteria so that parameters of simulated full-plate data sets, with a small amount of simulated noise, were recovered with high precision. To help the minimizer, I scaled *C*(0) values by a factor of 10^5^ to make them closer to other parameter values. For each plate, I ran many fits with different initial parameters to try to find a global minimum (see Section 2.6). I set parameter bounds according to Table 1, and checked that best fits had no parameters at a boundary.

**Table 1:**
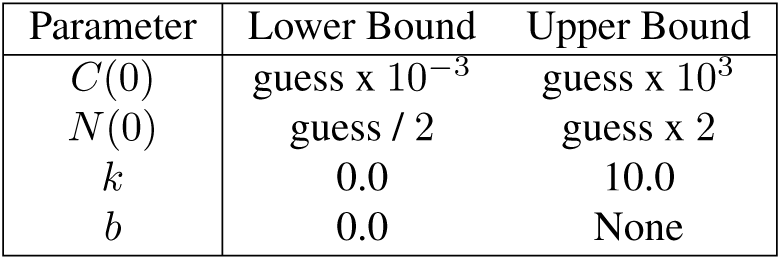
Parameter bounds for fitting the competition model to P15 and the Stripes and Filled plates. The same bounds on N(0) were used for both internal and edge cultures. Bounds on C(0) and N(0) were set using initial conditions: “guess” refers to the initial guess of each parameter (see Section 2.6).

Cultures at the edge of a plate gain an advantage from accessing a greater area of nutrients. I corrected for this using separate parameters, *N*_*I*_ (0) and *N*_*E*_(0), for the initial amount of nutrients in internal and edge cultures. In rate equations involving edge cultures, I scaled edge culture nutrient amount by the ratio *N*_*I*_ (0)=*N*_*E*_(0). The physical interpretation of this correction is that edge cultures have an extra supply of nutrients that can diffuse instantly into the reaction volume. This treatment increased the closeness of fit to cultures one row or column inside the edge, and to internal cultures overall (see Table 4 in Section 3.1.2). Cell density measurements from edge cultures contain extra noise due to reflections from plate walls (Lawless et al., 2010). For the competition model, unlike the logistic model, these cannot simply be discarded prior to fitting. Instead, I collectively fit to all cultures and selected best fits based on only the closeness of fit to internal cultures.

#### 2.4.3 Fitting the logistic model

Fitting the mass action logistic model requires a culture level *N*(0), creating 383 extra parameters. The QFA R package (Lawless *et al*., 2016) can fit the standard logistic model, and has heuristic checks to correct a confounding of *r* and *K* parameters that occurs when slow-growing cultures are dominated by noise. I did not have time to implement these checks for the mass action logistic model, so I instead fit using the QFA R package. This is not equivalent to the competition model fit because the QFA R package uses a culture level *C*(0), i.e., there is no collective fitting. This makes 1152 total parameters, rather than the 769 that we would have using the mass action logistic model. It is, nevertheless, useful to compare fitting of the competition model with the standard logistic model from the QFA R package, because the latter has been used in previous QFA papers, including Addinall *et al*. (2011) from which we take P15. In contrast to the competition model, noisy data from edge cultures can be discarded prior to fitting the logistic model. To remove bias in objective function comparison, I converted estimated logistic model parameters to competition model parameters (see Section 2.5) and reevaluated using CANS.

#### 2.4.4 Data visualisation

I used the Python package matplotlib to create plotting functions in CANS to visualise fits and simulations of QFA time-courses and to compare the ranking of fitness estimates.

### 2.5 Parameter conversion

In the independent limit (*k* = 0), it is possible to equate *C* of the competition and logistic model. Parameters can then be converted as follows:

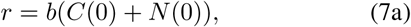

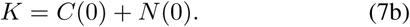

The competition model assumes that all nutrients are converted to cells, implying that all cultures starting with the same amount of nutrients reach the same final amount of cells. As final cell amounts generally differ, it is, therefore, necessary to allow *N*(0) to vary for each culture when fitting the mass action logistic model. This is not realistic; the competition model already assumes *k* > 0. It is, nevertheless, possible to disregard our interpretation of the competition model, fit the mass action logistic model, and convert parameters to those of the standard logistic model. This is illustrated in Figure 4a showing a single culture from a 16×24 format plate. This culture grew faster than its neighbours (not shown) and reached a higher final cell amount. Notice that estimated *N*(0) is approximately equal to final cells. The conversion equations can also be used to return corrected *r* and *K*, for growth in independent conditions, after first fitting the full competition model. This is illustrated in Figure 4b, in a fit to the same culture, where dashed lines represent the corrected fit. Notice that corrected values of *r* and *K* differ from those in Figure 4a.

It is easy to compare competition model *b* with logistic model parameters, because *C*(0) and *N*(0) are fit collectively for each plate. This leads to *b*∞*r*∞MDR (see Equations 2 and 6). Note also that all cultures have the same *K* and MDP. *b* is therefore equivalent to several common QFA fitness measures: *r*, MDR, and MDR MDP (Addinall *et al*., 2011). This makes *b* a very convenient fitness measure; we need not convert to logistic model parameters to compare the fitness rankings of cultures on the same plate. To compare fitness rankings between different plates we can of course use *b*. This is not, however, equivalent to comparing *r* or MDR as different plates may have different *C*(0) and *N*(0).

**Figure 4:**
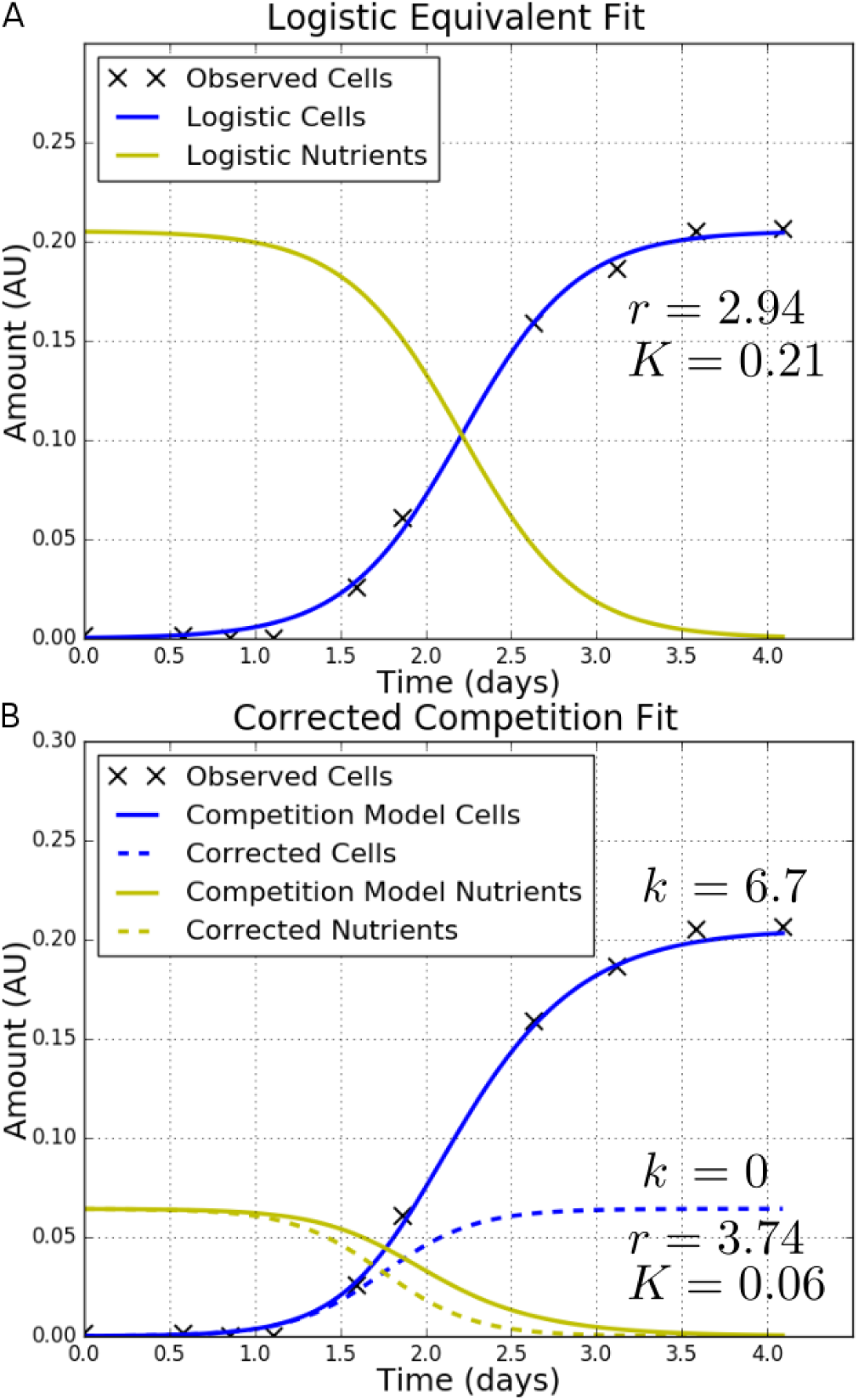
Using the competition model to return logistic model parameters. **A**) Fit of the logistic model displaying estimated parameters. **B**) Fit of the competition model (solid; k = 6:7), and simulation of independent growth (dashed; k = 0) displaying corrected logistic model parameters. Both fits are to culture (R10, C3) of P15 which grew faster and reached a higher final cell density than its neighbours (not shown).

### 2.6 Determining initial parameters

Fitting the competition model to a full plate with 387 parameters is a formidable optimisation problem with potentially many local minima. Achieving a good fit therefore requires a good initial guess. To fit small simulated zones, I simply used many random parameter guesses. For a full plate, more sophisticated guessing methods are required because the chance of any random guess being close to optimal values diminishes. I developed the *Imaginary Neighbour Model* (Figure 5) for guessing competition model *b* and this allowed good fits to be made.

**Figure 5:**
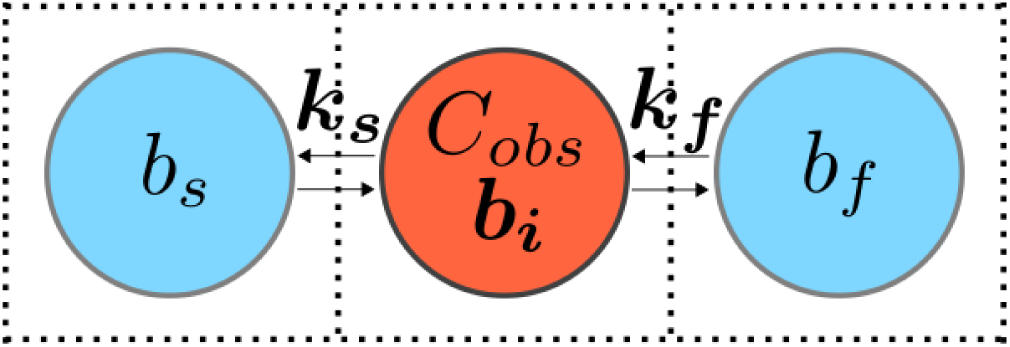
Schematic of the imaginary neighbour model. A real culture (red) with cell observations, C_obs_, is modelled growing alongside imagined fast and slow growing neighbours (blue) with growth constants b_f_ and b_s_ (for fast and slow). Rather than a single parameter, the model uses separate nutrient diffusion constants, k_f_ and k_s_, for the fast and slow growing neighbours. k_f_, k_s_, and the growth constant, b_i_, of the real culture are estimated by fitting the model to C_obs_ with all other parameters fixed. b_i_ are then taken as initial guesses for the competition model. Different numbers of each neighbour can be chosen to replicate different configurations of neighbours that might be present on a real plate.

### 2.6.1 Guessing initial amounts

To fit the imaginary neighbour model, it is first necessary to guess initial amounts. Recall from the competition model reaction equations (3 and 5) that nutrients can only diffuse or be converted to cells. Thus, assuming that reactions are nearly complete at the end of cell observations, and that *C*(0) ≪ *C*(∞), total initial nutrients, *N*_*Tot*_, can be estimated as,

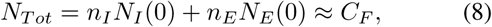

where *C*_*F*_ is total final cells, *n*_*I*_ and *n*_*E*_ are the numbers of internal and edge cultures, and *N*_*I*_ (0) and *N*_*E*_(0) are initial nutrient amounts for internal and edge cultures. Using this equation, and an estimate for the ratio of the area associated with edge cultures to the area associated with internal cultures, *A*_*r*_ = *A*_*E*_=*A*_*I*_ = *N*_*E*_(0)=*N*_*I*_ (0), I guessed *N*_*I*_ (0) and *N*_*E*_(0) as follows:

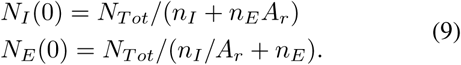

When *A*_*r*_ = 1, *N*_*I*_ (0) = *N*_*E*_(0) and we loose a parameter. I estimated *A*_*r*_ = 1:5 for plates in this paper.

When using dilute cultures, *C*(0) cannot be detected. I used a range of values in logspace and made a fit for each, rather than estimate a single initial guesses. The range was chosen to encompass uncertainty in *C*(0) for the given experiment.

### 2.6.2 Guessing *b*

From initial guesses of *N*(0) and *C*(0), I used the imaginary neighbour model (Figure 5) to quickly fit individual cultures and obtain initial guesses of *b*_*i*_. This model is based on the reaction and rate equations of the competition model (3–5), but tries to replicate the diffusion of nutrients into and out of a culture using imaginary fast and slow growing neighbours. These have growth constants *b*_*f*_ and *b*_*s*_, and separate nutrient diffusion constants *k*_*f*_ and *k*_*s*_. To fit, I fixed *C*(0) and *N*(0) as the initial guesses, *b*_*f*_ at a high value, and *b*_*s*_ = 0. I allowed *b*_*i*_, *k*_*f*_, and *k*_*s*_ to vary from zero to infinity. I used initial values of zero for *k*_*f*_ and *k*_*s*_. For each set of parameter guesses, I ran fits with initial values of *b*_*i*_ over the range (35, 40, …, 100), with *b*_*f*_ fixed at 1.5 times this value. I determined the number of imaginary neighbours, such that the culture with the highest observed final cell density had enough slow growing neighbours to provide all of the nutrients necessary to reach this value. I solved the imaginary neighbour model using SciPy’s integrate.odeint and fit using a gradient method as in Section 2.4.2. Fits of the imaginary neighbour model take several minutes compared to several hours for the the competition model.

### 2.6.3 Guessing *k*

The last parameter left to guess is nutrient diffusion constant *k*. I observed that simulations of the competition model using *b*_*i*_ drawn from normal distributions have linear relationships between variance in final cells, 
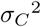
 and *k*. Setting other parameters as the guesses above, I simulated the competition model over a range of *k* and fit a straight line to 
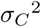
 vs *k* using simple linear regression. I then took measured 
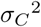
 and used the line to predict *k*.

## 3 RESULTS

### 3.1 Model comparison using P15

#### 3.1.1 Quality of fit

I compared the competition and logistic models by fitting both to P15 (data described in Section 2.2). For the competition model, I used five values of C(0) over the range *N*(0) 10 ^5^ < *C*(0) < *N*(0) 10 ^3^ to make initial parameter guesses as described in Section 2.6. This generated 70 initial parameter sets which I used in 70 separate fits. Cell density estimates fit data closely (Figure 6). A high nutrient diffusion constant, *k*, is estimated, such that nutrients diffuse readily and nutrient timecourses are similar across local areas of the plate. Estimated parameters for the two closest fits are show in Table 2. The fits agree better for *N*_*I*_ (0) and *N*_*E*_(0) than for *C*(0) and *k*. It appears that the gradient method is not finding a global minimum. However, b_i_ estimates are correlated with Spearman’s rank correlation coefficient, ρ_*S*_ = 0:989, and have average mean absolute deviation, MAD = 1:56. The mean value of b for the best fit is 44.4, so agreement between *b*_*i*_ are good. This is important because b is to be used as a fitness estimate to compare with the logistic model.

**Table 2:** Estimated parameters for the two best competition model fits to P15. Spearman’s rank correlation coefficient (ρ_S_) between b estimates is 0.989. Mean absolute deviation (MAD) between b estimates is 1.56. Mean b for the best fit is 44.4. Obj. is the total objective function value for internal cultures (smaller values are better).

| Fit | $C(0)$ | $N_I(0)$ | $N_E(0)$ | $k$ | Obj. |
| --- | --- | --- | --- | --- | --- |
| 1st | $9.1 \times 10^{-5}$ | 0.064 | 0.090 | 6.7 | 0.194 |
| 2nd | $13.9 \times 10^{-5}$ | 0.062 | 0.097 | 8.3 | 0.196 |

**Figure 6:**
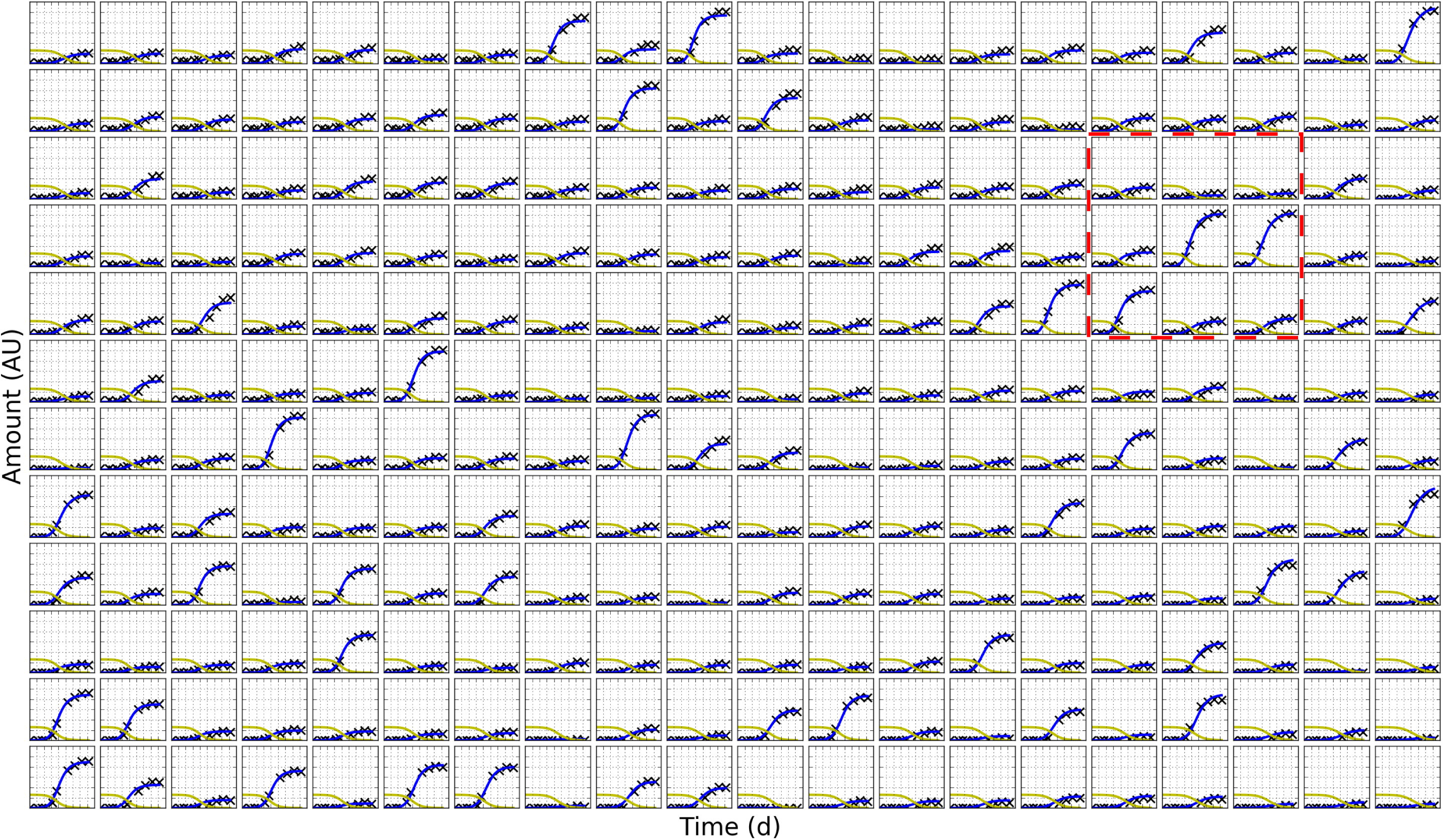
Fit of the competition model to P15. Data are for a 16×24 format plate (P15) with background mutation cdc13-1 incubated at 27°C. The plate contains 6 repeats of 50 genetic strains arranged randomly across internal cultures. The outer two rows and columns have been removed from the image to better fit the page. Model output for state variable, cell population size (blue curve), is fit to observed data (black crosses). Model predictions for unobserved variable (nutrient amount) are also plotted (yellow). The boxed zone is plotted at a larger size in Figure 7.

The boxed 3×3 zone in Figure 6 is replotted in Figure 7 with fits of the competition model (solid blue and yellow), initial guess (dashed blue and yellow), and logistic model (solid red). The zone contains more fast growing cultures than is typical for the plate, so competition effects might be greater than average. The fit of the initial guess is typical for the plate: timecourses for some cultures, such as the bottom-left, are poor. Nevertheless, the guess fulfils its purpose by allowing a good fit of the competition model to be made. Objective function values for the logistic and competition model are similar for most cultures in the zone. For the centre and centre-right fast growing cultures, however, the competition model has lower objective function values and fits are much closer. The total objective function value for the zone is lower for the competition model: 44.06 vs 68.38 (values scaled by 10^4^). This is not typical for the plate: objective function values are on average slightly better for the logistic model (see Table 3). Recall that only internal objective function values are used to assess the fit. Overall, the competition model performed well considering that it used far fewer parameters (387 vs 1152).

**Figure 7:**
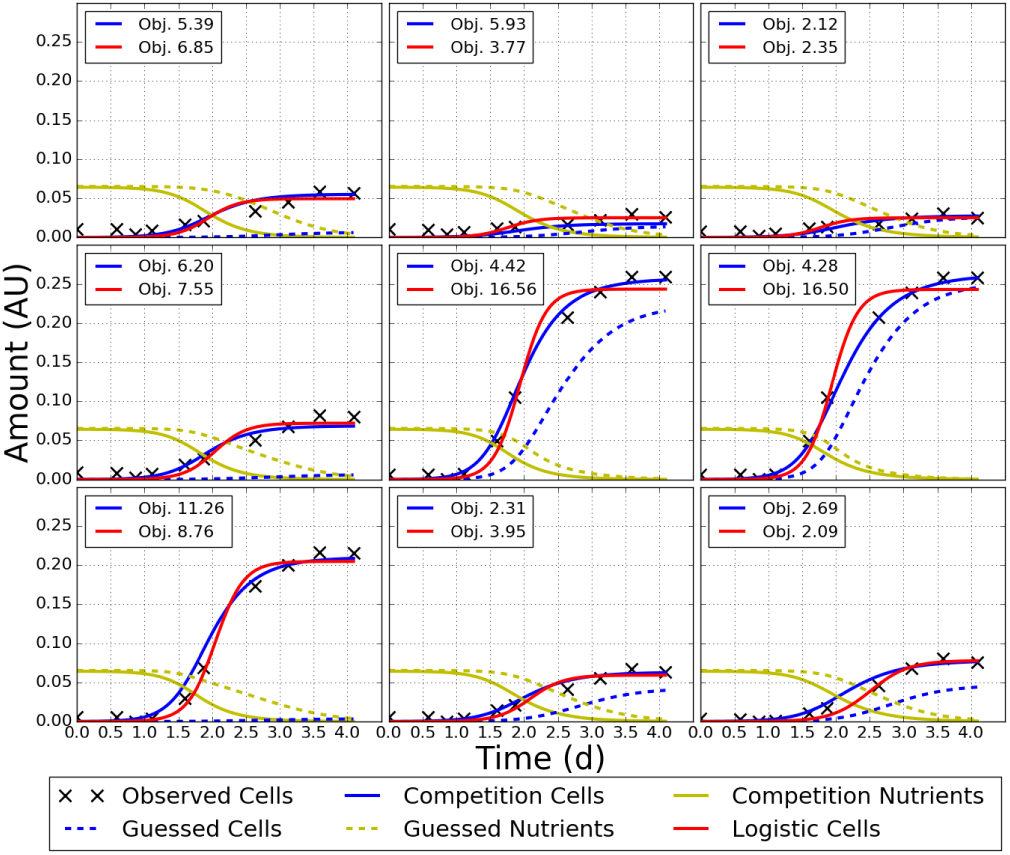
A 3×3 zone of P15 showing fits of the competition and logistic models. The zone has top-left coordinates (5, 18) and is boxed in red in Figure 6. This is a zoom on a whole plate fit, not just a fit to the zone. Fits are for the competition model (blue and yellow solid); logistic model (solid red), and initial guess (blue and yellow dashed). Objective function values (Obj.), scaled by 10^4^, are displayed for each culture (smaller values are better). Total objective function for the competition model is 44.06, for the logistic model 68.38.

**Table 3:** Objective function values for competition and logistic model fits to P15. “Internal” is the total objective function value for cultures not at an edge and is used to select the best fit. “All” is the total objective function value for all cultures on the plate. Smaller values are better.

| Cultures | Competition | Logistic |
| --- | --- | --- |
| Internal | 0.194 | 0.155 |
| All | 0.465 | 0.345 |

#### 3.1.2 Evaluating the treatment of boundaries

To evaluate how I treat boundaries, I again fit the competition model to P15, this time without a separate parameter for the initial amount of nutrients in edge cultures. Recall that edge cultures must be included when fitting the competition model, despite being discarded from results due to noise. The quality of fit appears similar for both the one and two *N*(0) models (Figure 8); the average objective function values for the entire plate are similar but slightly better for the two *N*(0) model (Table 4). Surprisingly, objective function values for edge cultures are worse for this model. Nevertheless, the fit is improved for cultures next to an edge, and, importantly, for internal cultures overall. The two *N*(0) model takes a deficit from edge cultures to improve the overall fit.

**Figure 8:**
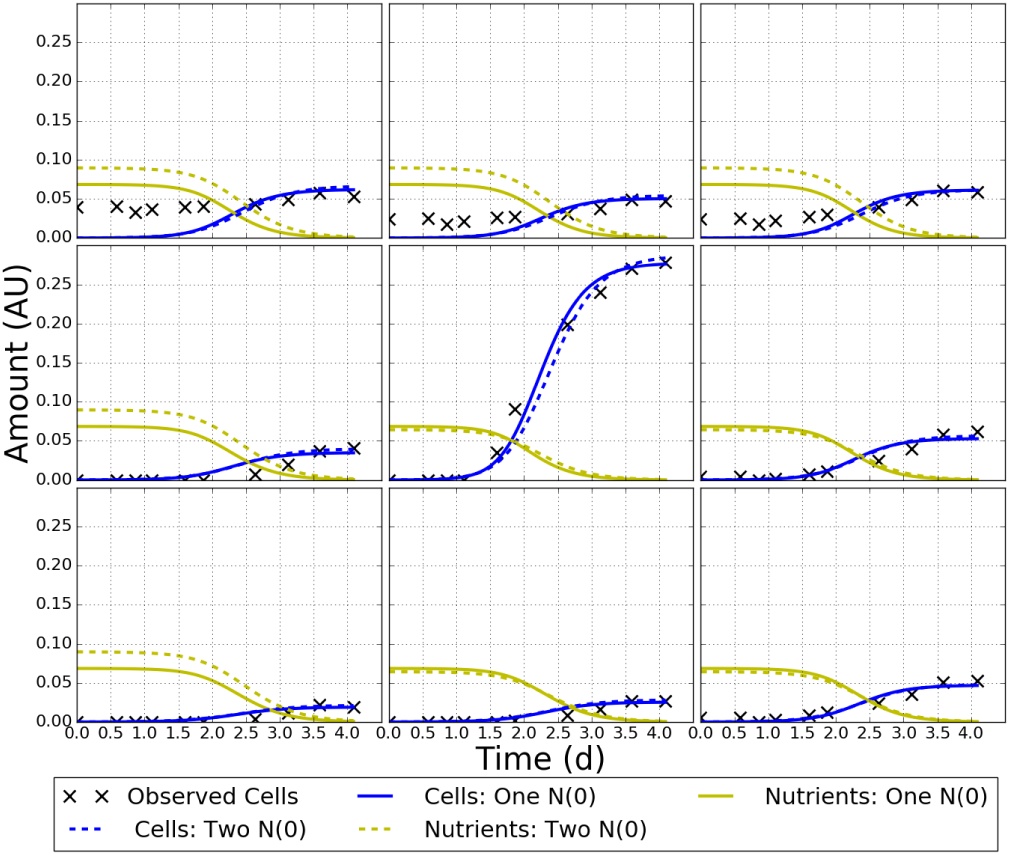
Fits of one and two initial nutrient parameter competition models to P15. The top left corner of a 16×24 QFA plate (P15) fit with two versions of the competition model: the first (solid) with a single initial nutrient amount for all cultures, the second (dashed) with a separate initial nutrient amount for edge cultures.

**Table 4:** Average objective function values for one and two N(0) parameter competition models fit to P15. Lower values are better. Averages are for cultures belonging to the areas indicated in the column “Cultures”. “Next to edge” refers to cultures one in from the edge. “Internal” refers to all cultures but the edge. Values have been scaled by 10^4^.

| Cultures | One $N(0)$ | Two $N(0)$ |
| --- | --- | --- |
| Edge | 35.9 | 36.5 |
| Next to edge | 9.54 | 7.98 |
| Internal | 6.67 | 6.30 |
| All | 12.4 | 12.2 |

#### 3.1.3 Correlation of fitness estimates

To compare fitness estimates between models, I converted competition model *b* to logistic model *r* using (6). I plotted correlation between *r* estimates for each culture (black) and between median *r* estimates for each deletion (red) (Figure 9). For individual cultures, the competition model distribution is unimodal, the logistic model distribution bimodal with minimum at *r*≈4. There are two distinct correlated groups: the first with lower *r* and gradient close to one, the second with higher *r* and a steeper gradient. Cultures from the middle of the competition model distribution, but not the tails, are split between groups. Competition model *r* was lower than logistic model *r* for almost all cultures. There are several outlying cultures (black) caused by failures of heuristic checks from the QFA R package. For the eight outliers on the left axis, checks set *r* = 0. The two outliers with very high logistic *r* are the deletions *rad50*Δ and *est1*Δ. They have escaped heuristic checks and exhibit a confounding effect between *r* and *K*.

**Figure 9:**
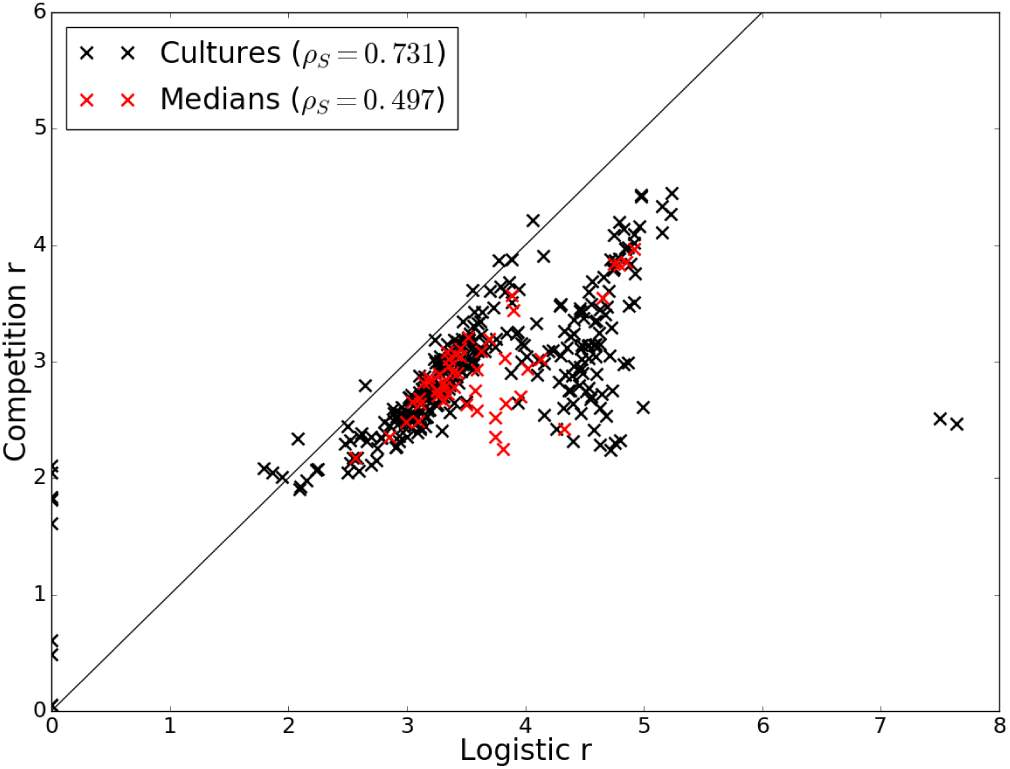
Correlation between logistic and competition model r for P15. Correlations are between estimates for each culture (black) and between median estimates (six repeats) for each deletion (red). Spearman’s rank correlation coefficient, ρ_s_, is displayed for each distribution. The line y = x is also plotted.

In the distribution of medians (red), most deletions fall inside the denser group of cultures with lower logistic *r*. Several deletions, with high competition model *r*, fall inside the second group (top-right). A significant number of deletions, however, lie in the region between the two groups, meaning that repeats of these cultures are split between groups. This may also be true for other deletions. Correlation, measured by Spearman’*s* rank correlation coefficient, ρ_S_, is lower between medians (0.497) than between cultures (0.731).

#### 3.1.4 Comparison of fitness rankings

I compared between models the ranking of strains by median fitness (Figure 10). Rankings of competition model *b*, *r*, and MDR are equivalent (see Section 2.5), so I compared b directly with logistic model *r* and *M D R*. The latter agree in the order of all but two deletions: *rad50*△ and *est1*△. These deletions each have an explained outlier with unrealistically high *r* and low *K* (Section 3.1.3). This appears to have been corrected for when *M D R* was calculated (2a), and *M D R* rank agrees much better with the competition model. Spearman’s Rho is therefore higher between competition *b* and logistic *M D R* (0.635) than between competition *b* and logistic *r* (0.497). Between models, there is good agreement in extreme positions, but middle positions are almost inverted. The positions of the neutral deletion, *his3*Δ (bold), disagree by 13 places.

**Figure 10:**
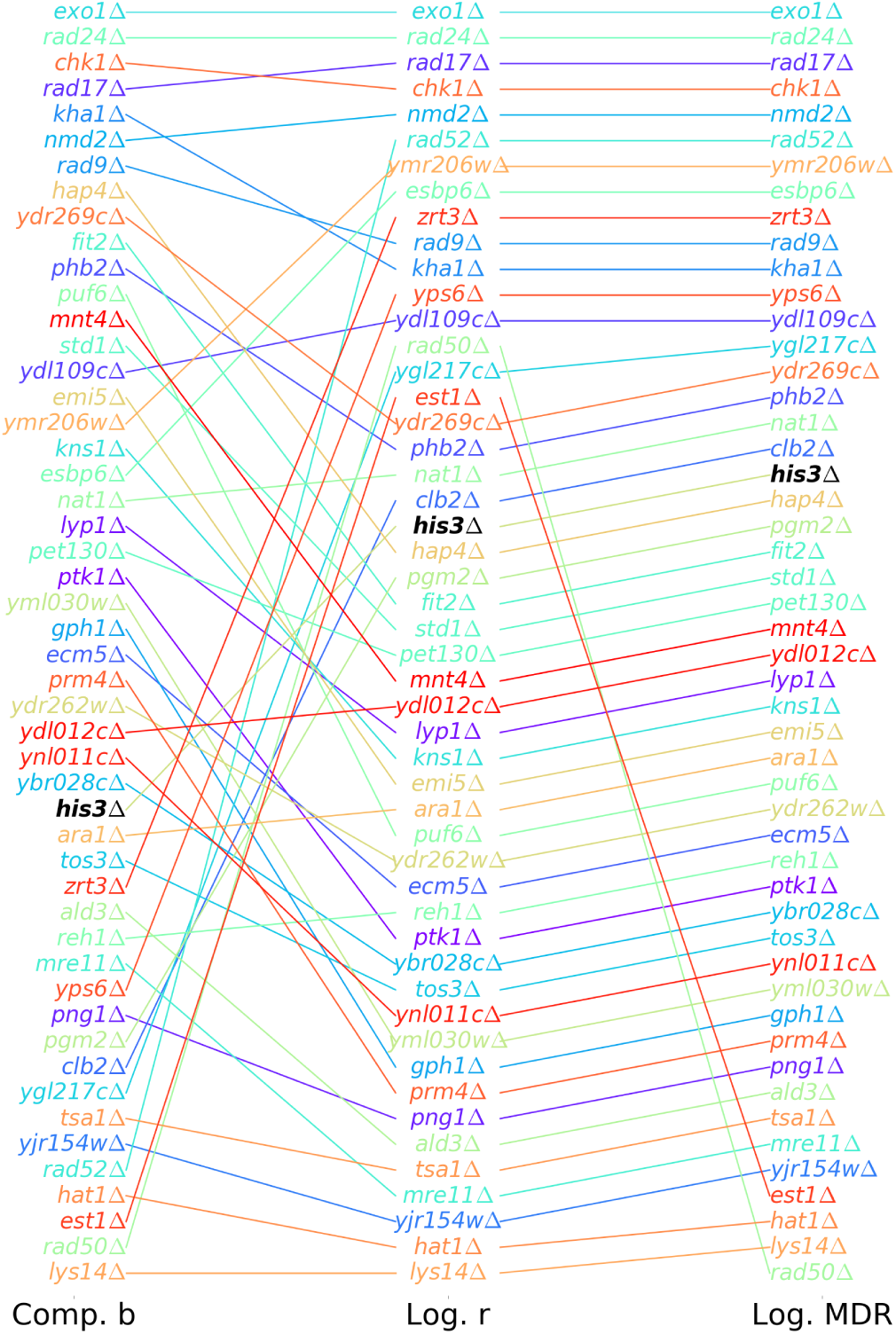
Comparison of competition and logistic fitness rankings for P15. Deletions, each with six repeats, are ranked according to median fitness, with the fittest strain at the top. Spearman’s Rho is 0.497 between competition b and logistic r, and 0.635 between competition b and logistic MDR. **his3Δ** is a neutral deletion with 14 repeats.

#### 3.1.5 Precision of fitness estimates

Using coefficient of variation (COV), I compared the precision of competition and logistic model fitness estimates in repeats of each deletion (Figure 11). COV of competition *b*, *r*, and MDR is equivalent. I chose to compare to logistic *r*, rather than MDR, because *r* and *b* are fixed properties of a strain, whereas MDR is dependent on initial conditions (see Equation 2a). This makes the former more useful for cross-plate comparison. Regardless, COVs for logistic *r* and MDR (not shown) are very similar. The competition model is more precise for 36 out of 50 deletions, but less precise for the 11 fastest growing (according to *b* ranking) strains. The precision of both estimates tends to decrease for slower growing strains.

**Figure 11:**
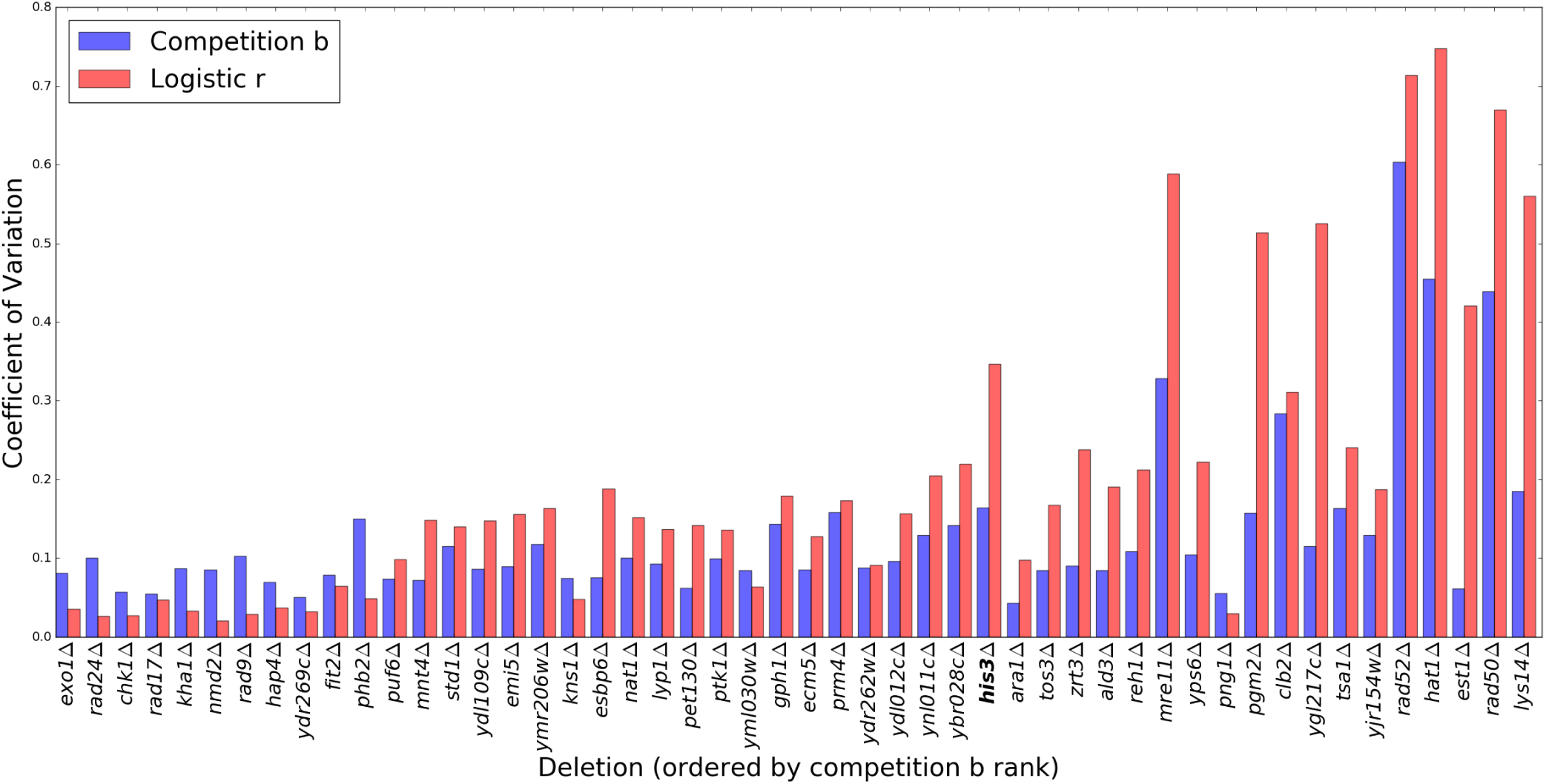
Coefficient of variation (COV) in P15 fitness estimates. Shown for each deletion are COVs in estimates of competition model b (blue) and logistic model r (red). COV in competition model b and r are equivalent so comparison is direct. Deletions are ordered left to right from highest to lowest b estimate. Each deletion has 6 repeats, except for the neutral deletion **his3Δ** which has 14. The competition model is more precise for 36 out of 50 strains, but less precise for the 11 fastest growing.

#### 3.2 Cross-plate validation

I conducted cross-plate calibration and validation of the competition model using the Stripes and Filled plates experiment, which is described in detail in Section 2.3. As for P15, I ran multiple fits to each plate (see Section 2.4.2). This time I used a wider range of *C*(0) guesses: 10 values over *N*(0)×10^−7^ < *C* < *N*(0) ×10 ^−1^ in logspace. I made this decision because of the higher inoculum densities in this experiment, and as a result of discussion with Herrmann and Lawless, who had suggested, based on recent work, that heterogeneity exists within individual cultures such that only a small number of cell lines contribute significantly to final populations. Strains in the Stripes plate are inoculated in the same locations on the Filled plate, and empty locations on the Stripes plate are inoculated with extra strains on the Filled plate. When fitting, I ignored data for empty cultures which is just noise. I took the estimated parameters for the Filled plate and simulated the Stripes plate by setting b to zero for empty locations. The simulated timecourses can be compared with the Stripes cell observations to validate the competition model (Figure 12); if the model corrects perfectly for differences between the plates, then simulated timecourses (blue dashes) should fit closely to Stripes data (blue crosses). I found that the competition model overcorrects: simulated cell densities are higher than observations across the plate. In contrast, the logistic model makes no adjustment between plates and systematically underestimates observed cell densities by a similar margin. There are some issues with the experiment. For instance, the top right culture in Figure 12 does not grow and might have been inoculated with dead cells. Nevertheless, these mistakes are unlikely to explain the systematic overestimation that is observed.

**Figure 12:**
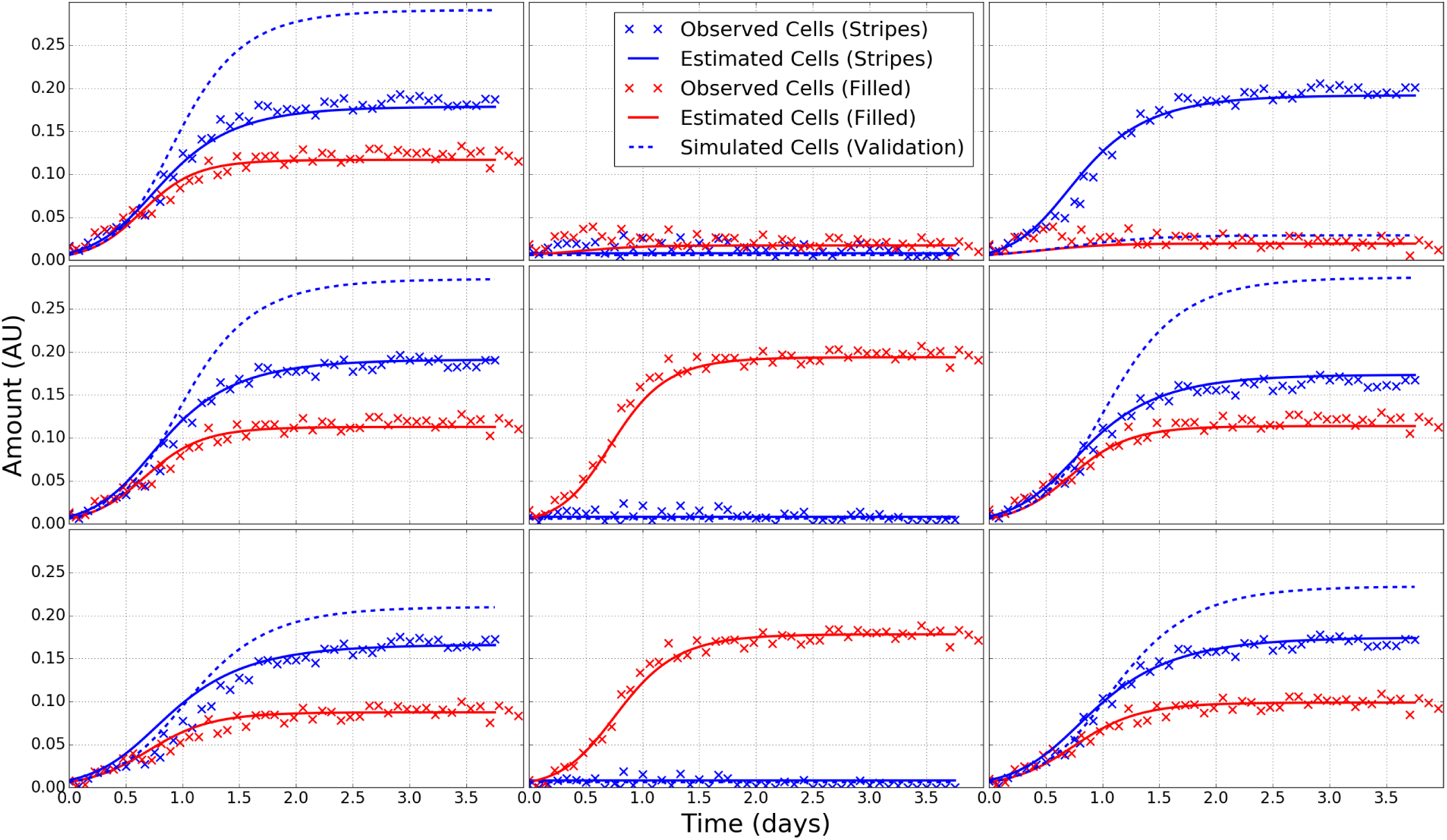
Cross-plate calibration and validation of the competition model. I fit the competition model to the 16×24 format Stripes and Filled plates (Section 2.3). The plot shows measured cells (crosses) and estimated cells (solid curves) from both plates for a 3×3 section with top left coordinates (R9, C10). I took the parameter estimates for the Filled plate (calibration), and set growth constants to zero for cultures in the empty columns of the Stripes plate (column two here). I then simulated using these parameters to produce the validation curves (dashed blue). If the competition model corrects for differences in growth between these experimental designs perfectly, the dashed blue curves should resemble the Stripes data (blue crosses) in columns one and three. The logistic model predicts no difference between plates.

I looked more closely at estimated parameters to investigate why the validation curves disagree with data (Table 5). All parameters disagreed significantly between plates. Both plates used the same inoculum density and formula of agar so I did not expect plate level parameters to differ so much. In particular, *k* is much higher for the Filled plate. Estimates of *C*(0) differed the least. The average *b* estimate among common cultures was significantly higher for the Stripes plate. In summary, the Stripes solution has faster growing cultures, fewer starting nutrients, and slower diffusion. Despite these differences, both solutions fit their data with similar closeness.

**Table 5:** Estimated parameter values for the best fit to the Stripes and Filled plates. The mean b is taken only for common cultures. Spearman’s rank correlation coefficient for b estimates between these cultures is 0.787.

| Plate | $C(0)$ | $N_I(0)$ | $N_E(0)$ | $k$ | $b$ (mean) |
| --- | --- | --- | --- | --- | --- |
| Stripes | $8.3 \times 10^{-3}$ | 0.085 | 0.096 | 1.9 | 39.5 |
| Filled | $6.2 \times 10^{-3}$ | 0.116 | 0.183 | 4.8 | 27.9 |

When examining fits to the same plate, agreement between parameter estimates is poor compared to P15; the five best fits to the Filled plate are only consistent for initial nutrient densities (Table 6). Despite this, all solutions overestimate growth when used to simulate the Stripes plate. Disagreement is greater still for the Stripes plate (not shown).

**Table 6:** Estimated parameter values for the best five competition model fits to the Filled plate. Estimates are ordered by whole-plate objective function value (Obj.). Lower values are better. Mean Absolute Deviation (MAD) in b estimates is calculated against the best fit for which μ_b_ = 27:9. Spearman’s Rho in b estimates is greater than 0.96 for all combinations of these fits.

| Obj. | $C(0)$ | $N_I(0)$ | $N_E(0)$ | $k$ | MAD $b$ |
| --- | --- | --- | --- | --- | --- |
| 0.76 | $6.2 \times 10^{-3}$ | 0.116 | 0.183 | 4.8 | - |
| 0.85 | $5.6 \times 10^{-3}$ | 0.119 | 0.175 | 3.1 | 0.77 |
| 0.88 | $5.0 \times 10^{-3}$ | 0.119 | 0.175 | 2.9 | 1.65 |
| 0.93 | $15.4 \times 10^{-3}$ | 0.114 | 0.171 | 4.7 | 11.1 |
| 0.97 | $3.2 \times 10^{-3}$ | 0.118 | 0.179 | 3.8 | 7.80 |

#### 3.3 Towards a genetic algorithm

In the interest of developing a quick method for estimating parameters while avoiding local minima, I investigated how well culture level parameters can be recovered when plate level parameters are fixed. The idea is to evolve candidate sets of plate level parameters and evaluate by fitting data using a gradient method. If culture level parameters can be recovered reliably, we might not have to worry about confounding between plate and culture level parameters when evolving candidates. I took parameter estimates from the fivebest fits for P15, and simulated full plate timecoureses at the observed times. I added a small amount of random noise to imitate real data. I then fixed plate level parameters to the true values and fit the competition model just for b. I set bounds of 0 < b < 200 and used 11 evenly spaced guesses of b in the range 
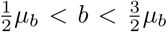
 where _b_ is the mean true b. These guesses were used in the imaginary neighbour model to generate 11 sets of initial parameters for the competition model for each simulation (Section 2.6.2). I solved all models using SciPy’s integrate.odeint, as I had not yet developed code to solve using libRoadRunner. Otherwise, I fit as for P15 (Section 2.4.2). *b* were recovered well even in the worst instance (Figure 13). In all instances, estimates were less accurate for slow growing cultures which are more affected by noise. For some simulations, different initial parameter sets all achieved a similar quality of fit. For others, however, certain guesses were much better, suggesting that multiple fits need to be run for each candidate.

**Figure 13:**
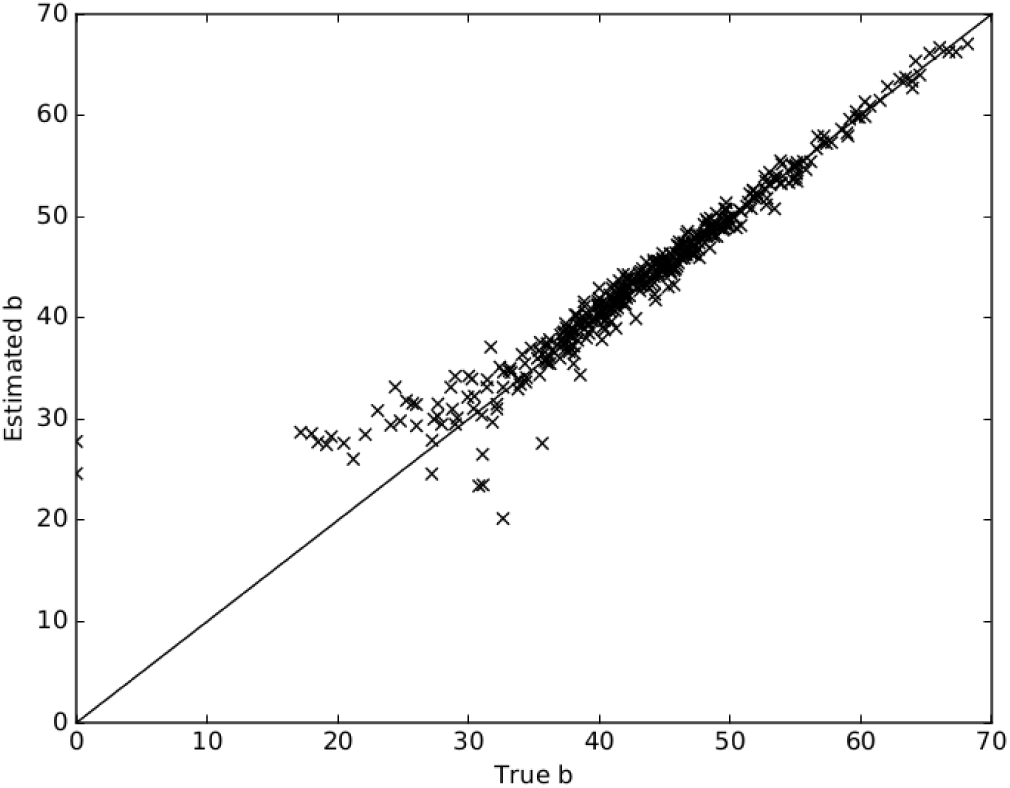
Recovery of true b values from a gradient fit with fixed plate level parameters. I simulated timecourses from the best five competition model fits to p15, added a small amount of noise, and used a gradient method to recover b given the true plate level parameters. This plot shows the worst recovery for the five sets of values.

### 4 DISCUSSION

On average, the competition model fit internal cultures of P15 worse than the logistic model: objective function value was 0.194 vs 0.155 (Table 3). However, for the two fastest growing cultures in a 3x3 zone studied in more detail, the competition model fit was better by far (Figure 7). I am yet to confirm that this trend carries across the plate, but it is likely that the logistic model performed better in regions where there were more slow growing cultures and less difference in growth between neighbours. The logistic model is more flexible when fitting slow growing cultures which are more affected by noise; it fits cultures independently with three culture level parameters, whereas the competition model fits cultures with three plate level parameters and only one culture level parameter. Logistic parameter estimates for slow growing cultures were less precise (Figure 11), and might be less accurate, despite the closer fit. Noise from slow growing cultures will also have affected competition model fits for fast growing cultures through plate level parameters and neighbour interactions.

Estimates from edge cultures are discarded due to noise from reflections off plate walls. Unlike the logistic model, the competition model must fit these cultures. For the fit of P15, edge cultures contributed 70% of error in objective function value despite making up only 20% of cultures (Tables 3 and 4). This will have affected all parameter estimates because cultures were fit collectively. However, it will have most greatly affected cultures neighbouring an edge. A two N(0) parameter model improved the fit of these cultures (Table 4) and might also have improved the reliability of estimated parameters. Considering the above effects and the fewer number of parameters used (387 vs 1152) the closeness of fit of the competition model is fairly good. Below I discuss differences in estimated parameters to discern whether one model made more reliable estimates.

For P15, the distribution of logistic model r estimates is bimodal, whereas that of the competition model is unimodal (Figure 9). It is possible that the true distribution is bimodal because Addinall *et al*. (2011) selected deletions suspected to have large positive and negative interactions with the query mutation. Nevertheless, repeats of the same strain are split between the gap in the logistic distribution; it appears that some deletions are acting as both enhancers and suppressors, but not as neutrals. It is possible that the split is an artefact of heuristic checks conducted by QFA R, or caused by a relative acceleration and deceleration of fast and slow growing cultures due to competition. If the latter case is true, the competition model might have corrected for it. It is important to investigate the cause of the bimodal distribution because it will affect the significance given to genetic interactions (see e.g. Addinall *et al*. (2011)).

Competition model fitness rankings of deletions on P15 are similar to the published logistic model rankings from Addinall *et al*. (2011) for the fastest and slowest growing strains (Figure 10). Both models also agree with rankings from independent spot tests for these strains (Maringele and Lydall, 2002; Zubko *et al*., 2004; Holstein *et al*., 2014; Foster *et al*., 2006,). Middle rankings, however, disagree between models, and Spearman’s Rho between *M D R* estimates is just 0.635. I investigated a disagreement between models by looking at plate images. *hap4*Δ is much healthier than *zrt3*Δ which agrees with the competition model and not the logistic model (Figure 10). This is promising for the competition model. However, I discuss below how inaccuracy in logistic estimates for the slower growing strain, *zrt3*Δ, may have been due to noise in the data rather than any fault of the model. Unfortunately, I lack independent data for middle strains to conclusively determine which model is more accurate. Collecting this data should be a priority for future studies.

The competition model was more precise for the majority of strains (36 out of 50), but less precise for the 11 fastest growing (see Figure 11). The fastest growing strains are not split across the gap in the logistic model distribution. Features of each model contribute to the differences in precision.

The logistic model fits cultures individually so it can be more precise for fast growing cultures which are less affected by noise. Conversely, the competition model fits cultures collectively, worsening the precision for fast growing cultures but improving the precise for slow growing cultures. If we believe that biological variation is low, it is reasonable to expect the better model to make more precise estimates for each strain. However, the higher precision for slow growing strains in the competition model could be entirely due to collective fitting rather than accuracy of the model. Similarly, the higher precision for fast growing strains in the logistic model could be due to a failure to account for competition. Although improved with the competition model, the precision of estimates for the slowest growing strains is still much lower than for the fastest, so the power to infer genetic reactions might not be dramatically improved.

Fitting the logistic model to slower growing cultures required heuristic checks to correct for confounding between *r* and *K*. The QFA R implementation appears to have some issues. The strains *est1*Δ and *rad50*Δ are very fast growing outliers in Figure 9. I visually inspected raw QFA images to confirm that these were weak growing strains. High *r* and low *K* were erroneously fit to both. This is corrected for when converting to *M D R*; rankings agree with the competition model and independent spot tests for rad50Δ (Zubko *et al*., 2004). *M D R* is similar to the fitness measure (*M D R*×*M D P*) used in the original analysis by Addinall *et al*. (2011) so these anomalies are unlikely to have affected their results. It is not clear how confounding is affecting other slow growing cultures. In some images it appears that encroachment of fast growing cultures into neighbours is affecting cell density estimates made by Colonyzer. If repeated, the plate from Addinall *et al*. (2011) should be run with a lower concentration of nutrients in the agar so that the stationary phase is reached before cultures start to merge.

In a validation experiment (Section 3.2), plates used a higher inoculum density reducing the effect of noise on slow growing cultures. The logistic model will not have required heuristic checks and the precision of competition estimates for fast growing strains should have improved. The experiment should therefore be a fairer comparison of the two models. Unfortunately, I struggled to find global minima with the competition model (Table 6) so the method of fitting needs to be improved before such a comparison can be made. Despite this, validation revealed issues with both models (Figure 12); the logistic model does not account for differences between plates at all and the competition model overcorrects. This was true for disparate solutions of the competition model so improving the method of fitting is unlikely redress the issue; it is likely that the competition model needs to be improved. With current fits, the logistic model undercorrects to a similar degree that the competition model overcorrects.

### 4.1 Future work

The distributions of logistic and competition *r* are very different (Figure 9) and can be used to compare models. First, P15 should be repeated using a higher inoculum density to eliminate the possibility that heuristic checks cause the split in the logistic distribution. If this is not the case, then strains could be grown in independent conditions - either isolated on solid agar (diffusion still an issue) or in liquid cultures - where the models are equivalent. We will be able to asses the quality of both models. The model whose QFA distribution is most similar to the independent distribution should be used to analyse QFA data.

I was unable to find global minima of the competition model using a gradient method (Sections 3.1 and 3.2). I found, however, that it is possible to reliably return *b*_*i*_ when plate level parameters are fixed at true values (Figure 13). It might therefore be possible to use a hierarchical genetic algorithm that evolves sets of plate level parameters, and evaluates using a gradient method to determine culture level parameters. Alternatively, a pure hierarchical genetic algorithm may work (i.e. where *b*_*i*_ are also evolved). A Bayesian hierarchical approach, similar to that of Heydari *et al*. (2016), could also return global minima, but would probably be much slower.

The systematic overcorrection seen in the validation experiment (Section 3.2) suggests that more fundamental improvements to the competition model are required. It would be informative to validate the independent limit to determine whether a mass action approximation is appropriate and whether it is correct to disregard the effect of metabolism on nutrient density. I suggest to validate first in liquid cultures, where the assumption of a well stirred mixture is more valid, and then attempt to validate for single cultures grown on agar, which more closely resembles QFA. The latter scenario has a lower dimensionality, and growth might be diffusion limited inside cultures, so a fractal kinetics model might be required (Kopelman, 1988; Savageau, 1995). Nutrients (sugars, nitrogen, etc.) in QFA agars are of a standard composition, designed to reduce the excess of any single nutrient (Addinall *et al*., 2011). It would be helpful to know and control the identity of the limiting nutrient using a different formula of agar. With nitrogen, rather than sugar, as the limiting nutrient, we are less likely to have to model metabolism.

Estimates of the nutrient diffusion constant *k* for P15 were fairly high (Table 2), such that nutrients diffused readily between neighbours and were nearly depleted when growth stopped (Figure 7). In reality, growth might become limited by the diffusion of nutrients through agar long before all nutrients are depleted, in which case nutrients are not well approximated as being evenly distributed across the spatial scales that we model. The Stripes fit might support this; cultures have access to more nutrients from empty neighbours and lower *k* and *N*(0) are estimated (Table 5). Using a finer grid could reduce the overcorrection seen in Figure 12. Reo and Korolev (2014) use the diffusion equation with Neumann and Dirichlet boundary conditions to simulate nutrient dependent growth of a single bacterial culture on a pertri dish in two-dimensions. They create a sink for nutrients from culture growth and equate the flux of nutrients through culture area with the rate of increase in culture size. They model culture area as varying and keep culture density constant. This model could be adapted for QFA by keeping culture area constant and allowing culture density to vary. A mass action kinetic model of reaction (3) could be used for culture growth and the nutrient sink. It might take too long to fit such a detailed model to a whole plate, but simulation could be used to determine an appropriate grid size for a mass action diffusion model.

If competition for nutrients is not responsible for the interaction between neighbours, for instance if growth stops before nutrients from neighbours can diffuse, then we could instead model signalling by ethanol poisoning (Fujita *et al*., 2006). Signal diffusion may be modelled as for nutrient diffusion (4,5) so code can be quickly adapted. If there is any combination of effects, e.g., competition, metabolism, signalling, overcrowding, or arrest contributing significantly to differences in the growth of cultures and the interaction between neighbours, then they will be difficult to separate when fitting a model. We may have to develop ways to calibrate effects in isolation and use this information when fitting to high-throughput data. Currently, QFA only observes cells.

### 4.2 Improvements and recommendations

Recent work by Herrmann and Lawless suggests that directly measurement of *C*(0) is unreliable due to heterogeneity between cells in the same inoculum; many cells do not grow and only the fastest growing cells contribute significantly to the final population. Modelling using a plate level *C*(0) seems inappropriate, but having to fit extra culture level parameters is undesirable. Instead, noting that only small amounts of nutrients are consumed when cultures are small, early time-points could be discarded, and *C*(0) could be measured after populations have undergone several divisions. Currently, QFA takes inocula from the stationary phase where there might be more heterogeneity (Bergkessel *et al*., 2016). It might be possible to reduce heterogeneity by taking inocula from the exponential growth phase. Alternatively, a higher inoculum density could be used to average out stochastic effects.

Noisy edge cultures, which must be included in fits of the competition model, contribute significantly to objective function values (see Table 4). Whereas these cultures reduce competition for the logistic model, noise would be better dealt with for the competition model if they were left empty. A different treatment of boundaries could also be used by modelling empty cultures outside edges rather than the approach in Section (2.4.2). It is quicker to fit to small zones of a plate, but these have a large proportion of edge cultures, and it is difficult to model the boundary. Growing smaller arrays in isolation would help to speed up the development process.

I thought of several improvements to the guessing method which I did not have time to implement. The current implementation of imaginary neighbour guessing (Section 2.6.2) uses a range of *b*_*f*_ values, but estimated *b*_*i*_ are chosen from sets of fits using the same value, i.e., *b*_*i*_ currently come from the same *b*_*f*_. In reality cultures have different arrangements of neighbours, so mixing of fits from different *b*_*f*_ should be allowed. Alternatively, *b*_*i*_ could be initialised by converting from logistic model estimates (Section 2.5). *k* is the last parameter to be guessed. Rather than using the linear relationship between variance in final cell measurement and *k*, which may not hold for large values, *k* could be guessed by fixing all other parameters as their guesses and fitting the competition model to data.

A dramatic change was made between the Stripes and Filled plates to ensure that competition was present. I could have first validated the model against a smaller change, by inoculating slower growing cultures in gaps rather than removing cultures completely. If the model behaves well in such an experiment, it may work for the majority of QFA experiments, which typically have smaller differences between cultures than the original validation experiment. We could also have calibrated and validated from the Stripes plate to the Filled plate, i.e., in the other direction, if the extra columns on the Filled plate were repeats of common cultures. A different validation experiment could try to induce changes in logistic model ranking by inoculating fast or slow growing cultures next to target strains. Hopefully competition estimates would show no change.

When inoculum density was low the logistic model struggled to deal with noise in slow growing cultures (Figure 11). It is desirable model nutrient dependent growth, so that information is shared between cultures using a plate level N(0), and to eliminate the dependence on heuristic checks.

